# MetaPhat: Detecting and decomposing multivariate associations from univariate genome-wide association statistics

**DOI:** 10.1101/661421

**Authors:** Jake Lin, Rubina Tabassum, Samuli Ripatti, Matti Pirinen

## Abstract

**Background:** Multivariate testing tools that integrate multiple genome-wide association studies (GWAS) have become important as the number of phenotypes gathered from study cohorts and biobanks has increased. While these tools have been shown to boost statistical power considerably over univariate tests, an important remaining challenge is to interpret which traits are driving the multivariate association and which traits are just passengers with minor contributions to the genotype-phenotypes association statistic.

**Results:** We introduce MetaPhat, a novel bioinformatics tool to conduct GWAS of multivariate and correlated traits using univariate summary results and to decompose multivariate associations into sets of central traits based on intuitive trace plots that visualize Bayesian Information Criterion (BIC) and P-value statistics of multivariate association models. We validate MetaPhat with Global Lipids Genetics Consortium GWAS results, and we apply MetaPhat to univariate GWAS results for 21 heritable and correlated polyunsaturated lipid species from 2,045 Finnish samples, detecting 7 independent loci associated with a cluster of lipid species. In most cases, we are able to decompose these multivariate associations to only 3-5 central traits out of all 21 traits included in the analyses. We release MetaPhat as an open source tool written in Python with built-in support for multi-processing, quality control, clumping and intuitive visualizations using the R software.

**Conclusions:** MetaPhat efficiently decomposes associations between multivariate phenotypes and genetic variants into smaller sets of central traits and improves the interpretation and specificity of genome-phenome associations.

MetaPhat is available under the MIT license at: https://sourceforge.net/projects/meta-pheno-association-tracer

## Introduction

Genome-wide association studies (GWAS) of common diseases and complex traits in large population cohorts have linked thousands of genetic variants to individual phenotypes. In emerging biobank studies, as well as in some disease specific collections focusing on, for example, Type 2 diabetes (T2D) (Mahajan et al., 2018) or coronary artery disease (CAD) (Ripatti et al., 2016), multiple related quantitative traits are simultaneously available for genetic association studies. The statistical power in these discovery efforts can be boosted considerably by multivariate tests which have become more practical through recent implementations that require only univariate summary statistics, such as METASOFT (Han et al., 2011), MultiPhen (O’Reilly et al., 2012), PLINK (Purcell et al., 2007), TATES (van der Sluis et al., 2013), CONFIT (Gai et al., 2018), MTAG (Turley et al., 2018), MTAR (Guo et al., 2019) and metaCCA (Cichonska et al., 2016). The merits of many of these methods are further discussed (Chung et al., 2019). Concretely, canonical correlation analysis (CCA) (Hotelling, 1936) is the direct extension of the correlation coefficient to identify linear associations between two sets of variables, and it has been successfully applied also to GWAS (Inouye et al., 2012). Moreover, metaCCA extended CCA to work directly from GWAS summary statistics (effect size estimates and standard errors) of related traits and studies. However, a remaining challenge is to interpret which traits are driving the multivariate association and which traits are just passengers contributing little to the association statistic. To address this important question, we introduce MetaPhat (Meta-Phenotype Association Tracer), a novel method to efficiently and systematically

1. Identify and annotate significant variants via multivariate GWAS from univariate summary statistics using metaCCA,
2. Perform decomposition by systematically tracing the traits of highest and lowest statistical importance to identify subsets of central traits at each associated variant,
3. Plot the traces of trait decompositions and cluster the variants based on the ranking of the importance of traits.

## Materials and Method

### Workflow

Open source MetaPhat software is written in Python with plotting scripts written in R. MetaPhat requires as input a set of related GWAS summary statistics from correlated traits. The program implements efficient multi-trait genome-wide association testing, identification of significant associations and systematic tracing of trait subsets to identify the central traits that consist of a statistically optimal set of traits together with a set of driver traits. A workflow is shown in Figure 1. In steps 1-3, genome-wide significant variants (P < 5e-8, the established genome-wide threshold in the field (Pe’er et al., 2008)) are identified and are clumped into independent groups that are subsequently represented by the lead variant of each group (i.e. the variant with the smallest P-value). By default, two lead variants are defined as independent if their distance is higher than 1 million bases. At step 4, we do the decompositions of multivariate association by starting from model with all K traits and removing one trait at a time until only one trait remains. We proceed via two different strategies that we name as the *highest trace* and the *lowest trace*. More specifically, starting from the model with all K traits, we test all unique combinations of (K−1) traits to find the subset with the highest CCA statistic (lowest P-value) that we assign to the highest trace, and the subset with the lowest CCA statistic (highest P-value) that we assign to the lowest trace. We continue both traces iteratively until only a single trait remains, by always choosing the subset with the highest CCA statistic on the highest trace and the subset with the lowest CCA statistic on the lowest trace. Altogether we evaluate K^2^ subsets out of all possible 2^K^ subsets while building these two traces. Base pair distances, GWAS P-value thresholds and other program parameters can be updated using command-line arguments.

**Figure 1:**
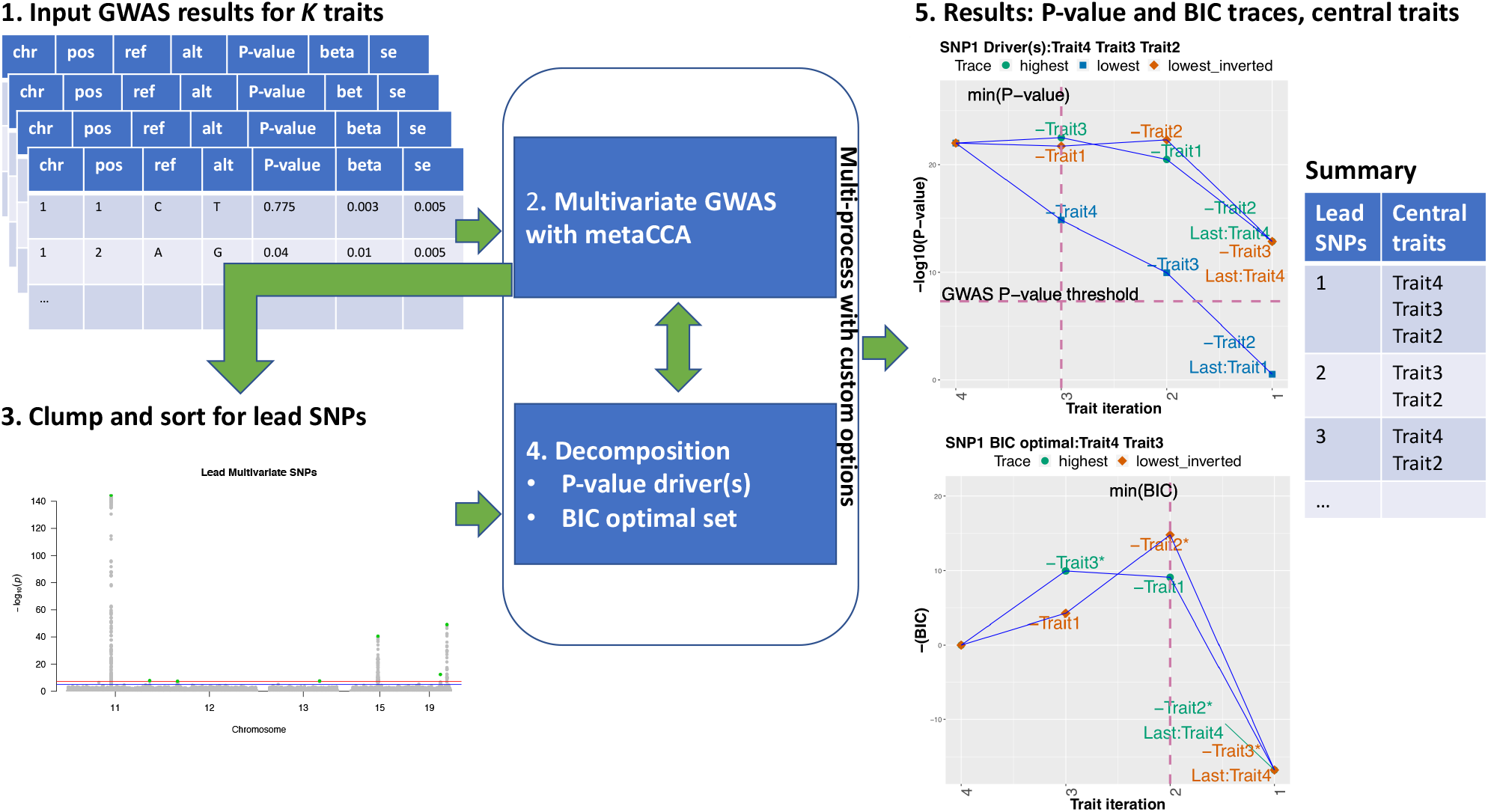
MetaPhat workflow. 1. GWAS results for K traits are accepted as input. 2. After quality control and filtering, multivariate GWAS is performed on the full model with all *K* traits using metaCCA via efficient multi-processing and chunking to reduce computation time. 3. Lead SNPs are detected and sorted based on the leading canonical correlation/P-value and then clumped based on a user-specified window size. Custom variants can be added. 4. Decomposition of chosen variants is performed through highest and lowest traces to find an optimal subset with a minimum BIC and driver traits based on the established P-value threshold. 5. MetaPhat results include trace plots for P-values and BIC, variant clusters based on trait importance and summary table listing central traits (union of drivers and optimal subset).

We use the two traces to identify central traits that are primarily responsible for the association with the variant as explained next.

### Evaluating models

We use two quantities to evaluate models: CCA P-values and Bayesian Information Criterion (BIC (Schwarz, 1978)). P-values allow comparing each association to the established “genome-wide significance threshold” of 5e-8 (Pe’er et al., 2008). By using the lowest trace we can identify those traits without which the multivariate P-value is no longer genome-wide significant by simply collecting the traits that have been removed from the full model when the P-value on the lowest trace is first time above 5e-8. We call these traits the **driver traits** as they drive the association in the sense that without them the association does not anymore reach genome-wide significance, and hence would not had been reported as a discovery in a GWAS. This definition of driver traits is based on a fixed P-value threshold, that is an established practice in the field, but does not claim any statistical optimality properties in terms of model comparison. Hence, to more rigorously compare models with different dimensionalities, we use BIC, that approximates the negative marginal likelihood of the model, and thus penalizes for the model dimension (Schwarz, 1978). A lower BIC value suggests a statistically better description of the data. Hence a subset of traits with minimum BIC would be the model of choice. We define the **optimal subset** as the subset with the lowest BIC among all subsets on the highest trace and all subsets on the inverted lowest trace. The inverted lowest trace aggregates the traits that have been dropped on the lowest trace, and, in particular, includes the set of the driver traits as one of its subsets. Subsequently, we define the **central traits** as the union of traits from the drivers and optimal BIC subset.

### Computing P-values and BIC from GWAS summary statistics

metaCCA outputs the first canonical correlation *r*_1_ between the genetic variant *x* and the set of *k* traits *y*_1_,…,*y*_k_ and computes the corresponding P-value (Cichonska et al., 2016). In this case, the first canonical correlation *r*_1_ equals to the maximum correlation between the variant and any linear combination of the traits, and hence equals to the square root of the variance explained *R*^2^ from the linear regression of *x* on *y*_1_,…,*y*_k_. In general, the expression for BIC is

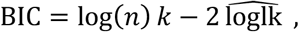

where *n* is the sample size, *k* is the number of parameters (here traits) and 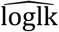 is the maximized log-likelihood. We next show how to use metaCCA output *r*_1_ to derive BIC from the maximized likelihood of the linear model written as a function of *R*^2^ = *r*_1_^2^.

Consider a linear model between a (mean-centered) variant *x* and (mean-centered) traits ***y*** = (*y*_1_,…,*y*_k_)^*T*^.

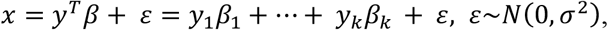

where we do not include the intercept parameter as its maximum likelihood estimate (MLE) is zero after mean-centering. Log-likelihood function is

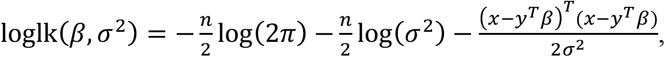

and MLEs are

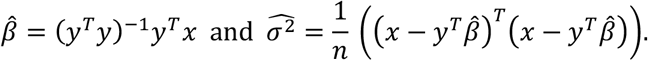

Thus, the log-likelihood at maximum is

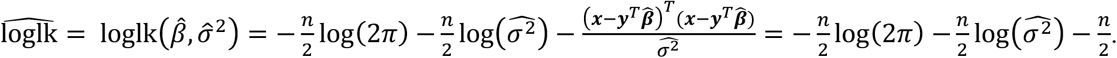

To connect this to the variance explained *R*^2^ note that

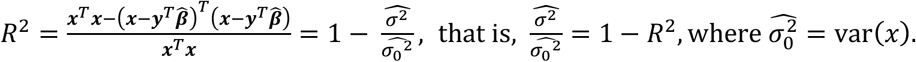

Hence, the logarithm of the likelihood ratio between the MLE and the null model can be written as

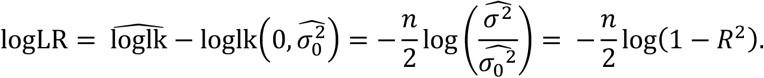

Hence, we have that, for an additive constant 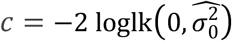,

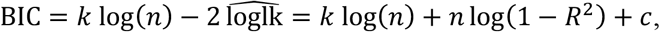

which is possible to compute directly from the metaCCA output for models with at least two traits up to an additive constant *c*. Since *c* does not depend on the model dimension, we can ignore it in the BIC calculation, when we are only interested in the differences in BIC between models.

Finally, for a single-trait model, *R*^*2*^ can be computed directly from the univariate GWAS summary statistics as

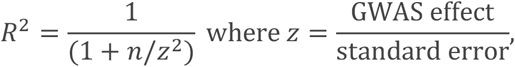

which can be plugged in the BIC formula above to yield BIC for the single-trait model.

### Implementation and output

MetaPhat is written in Python (compatible for 2.7 and 3+) and requires R (3.4+) for plotting. The command-line based program has been tested on multiple operating systems and cloud images. Library requirements and command options are further described in Supplement Table 1 and test data are accessible from the project page: https://sourceforge.net/projects/meta-pheno-association-tracer/

MetaPhat outputs tabular text files and several plots. A summary result file contains, for each chosen variant, the driver traits and the optimal subset with their P-value and BIC statistics. For each variant, trace plots using P-values and BIC are generated, showing the highest trace, the lowest trace and the inverted lowest trace. Additionally, the estimated phenotype correlation matrix and a clustered heatmap of trait importance for the variants are produced. Optionally, intermediate statistics during the decomposition can be plotted to get a more detailed view of the decomposition process.

## Materials

Our lipidomics data set consists of univariate GWAS results of 21 correlated lipid species with polyunsaturated fatty acids that were reported to exhibit high heritability (Tabassum et al., 2019) and showed high correlation (Figure S2). These results originate from 2,045 Finnish subjects with imputed genotypes available at ~8.5 million SNPs. The arbitrarily assigned lipid species identifiers along with their class names and fatty acid chemical properties are listed in Table 1A. To further validate MetaPhat, we processed summary statistics from four basic lipids conducted by the Global Lipids Genetics Consortium (GLGC) (Willer et al., 2013) and listed in Table 1B. With the GLGC data set our aim was to compare MetaPhat results with univariate results reported by GLGC for all variants reported to be significantly associated with two or more traits by GLGC.

**Table 1:** Lipid traits used in MetaPhat analysis. **A.** Polyunsaturated lipid species with acyl chains-C20:4 (14 lipids), C20:5 (3 lipids) and C22:6 (4 lipids) (Tabassum et al., 2019) measured for 2,045 individuals. After quality control (QC), a total of 8,576,290 variants were available for all 21 traits. **B.** Four well-studied lipids from GLGC (Willer et al., 2013). After quality control, a total of 2,267,285 variants were available for all four traits.

**A. Plasma lipidomics**
| Identifier | Lipid class | Lipid species | QC'd variants |
| --- | --- | --- | --- |
| CE14 | Cholesteryl ester | CE(20:4;0) | 8,711,715 |
| CE15 | Cholesteryl ester | CE(20:5;0) | 8,711,715 |
| CE17 | Cholesteryl ester | CE(22:6;0) | 8,711,665 |
| LPC8 | Lysophosphatidylcholines | LPC(20:4;0) | 8,710,151 |
| LPC9 | Lysophosphatidylcholines | LPC(22:6;0) | 8,694,250 |
| LPE5 | Lysophosphatidylethanolamine | LPE(20:4;0) | 8,710,162 |
| LPE6 | Lysophosphatidylethanolamine | LPE(22:6;0) | 8,711,037 |
| PC17 | Phosphatidylcholine | PC(16:0;0-20:4;0) | 8,711,715 |
| PC18 | Phosphatidylcholine | PC(16:0;0-20:5;0) | 8,711,533 |
| PC29 | Phosphatidylcholine | PC(17:0;0-20:4;0) | 8,704,982 |
| PC36 | Phosphatidylcholine | PC(18:0;0-20:4;0) | 8,711,715 |
| PC37 | Phosphatidylcholine | PC(18:0;0-20:5;0) | 8,751,062 |
| PC46 | Phosphatidylcholine | PC(18:1;0-20:4;0) | 8,711,715 |
| PC21 | Phosphatidylcholine | PC(16:0;0-22:6;0) | 8,711,715 |
| PCO7 | Phosphatidylcholine-ether | PC-O(16:0;0-20:4;0) | 8,711,715 |
| PCO23 | Phosphatidylcholine-ether | PC-O(18:0;0-20:4;0) | 8,711,560 |
| PCO29 | Phosphatidylcholine-ether | PC-O(18:1;0-20:4;0) | 8,710,292 |
| PE7 | Phosphatidylethanolamine | PE(18:0;0-20:4;0) | 8,707,361 |
| PEO3 | Phosphatidylethanolamine-ether | PE-O(16:1;0-20:4;0) | 8,706,846 |
| PEO11 | Phosphatidylethanolamine-ether | PE-O(18:2;0-20:4;0) | 8,693,147 |
| PI9 | Phosphatidylinositol | PI(18:0;0-20:4;0) | 8,711,715 |

**B. GLGC Lipids**
| Identifier | Lipid class | QC'd variants | Sample size |
| --- | --- | --- | --- |
| HDL | High density lipoprotein cholesterol | 2,343,025 | 95,129 |
| LDL | Low density lipoprotein cholesterol | 2,271,091 | 90,421 |
| TC | Total cholesterol | 2,341,292 | 95,537 |
| TG | Triglycerides | 2,286,633 | 91,598 |

## Results

Using the lipidomics data sets with GWAS summary statistics from the 21 polyunsaturated lipids, MetaPhat found 7 independent lead variants after clumping the 415 variants exceeding the standard GWAS P-value threshold of 5e-8 within a window of 1 Mb. Table 2 lists the results including which four of these seven variants were previously reported by GLGC as associated with at least one of the four basic lipids. The other three variants also have some nearby variants that have been reported in the GWAS catalog (Buniello et al., 2019). First, rs8736 in *MBOAT7* has been previously reported to be associated with human blood metabolites (Shin et al., 2014) as well as alcohol related cirrhosis of the liver (Buch et al., 2015). Second, variants in the region of rs146327691, near the *SLCO1A2* gene, have been previously reported for response to serum metabolites (Krumsiek et al., 2012) and, interestingly, also for response to statins (Ho et al., 2006, Carr et al., 2019). Lastly, variants in the region of rs188167837 have been previously identified to be associated with nasopharyngeal carcinoma (Su et al., 2013).

**Table 2:**
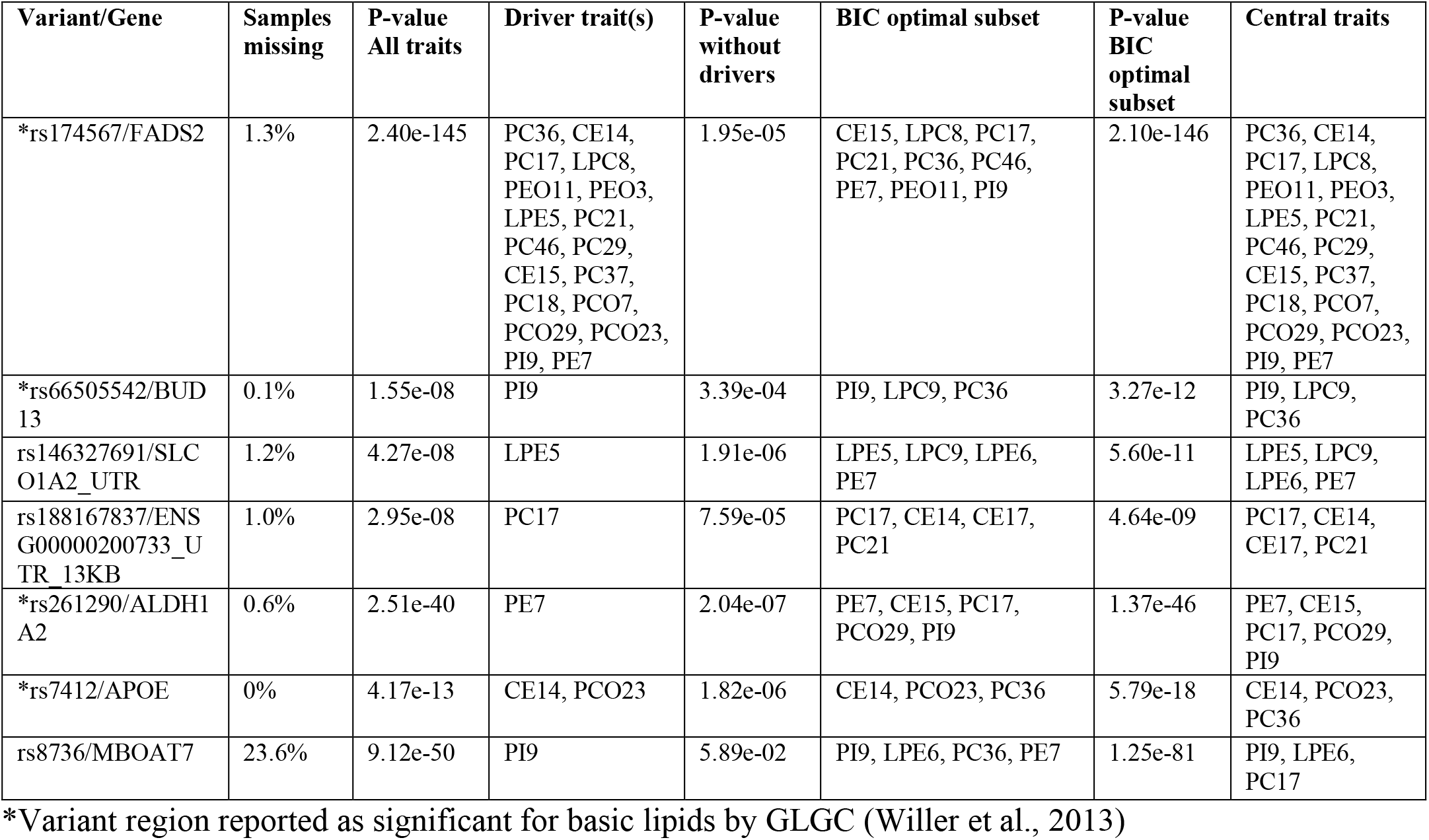
MetaPhat results of the 7 lead variants from the multivariate analyses of the lipidomics data. The lipid trait class names and acyl chain properties are listed in Table 1A.

In particular, let us consider in more detail rs7412 that is a missense variant in *APOE* gene and is known for its effect on LDL as reported, for example, in the GLGC analysis (Willer et al., 2013). With the lipidomics data this variant would not have been identified from any of the 21 univariate GWAS as the smallest univariate P-value was 1.1e-4. On contrary, the multivariate GWAS by MetaPhat clearly highlighted this variant associated with the multivariate lipidomics (P = 4.2e-13) and further determined that the association was driven by CE14 and PCO23 (P-value after excluding these driver traits is 1.8e-06). The BIC-optimal subset model for this variant extended the drivers by one additional trait and included CE14, PC36 and PCO23, which form the central traits. The trace plots for rs7412 are shown in Figure 2A (P-values for defining driver traits) and Figure 2B (BIC for defining optimal subset).

**Figure 2:**
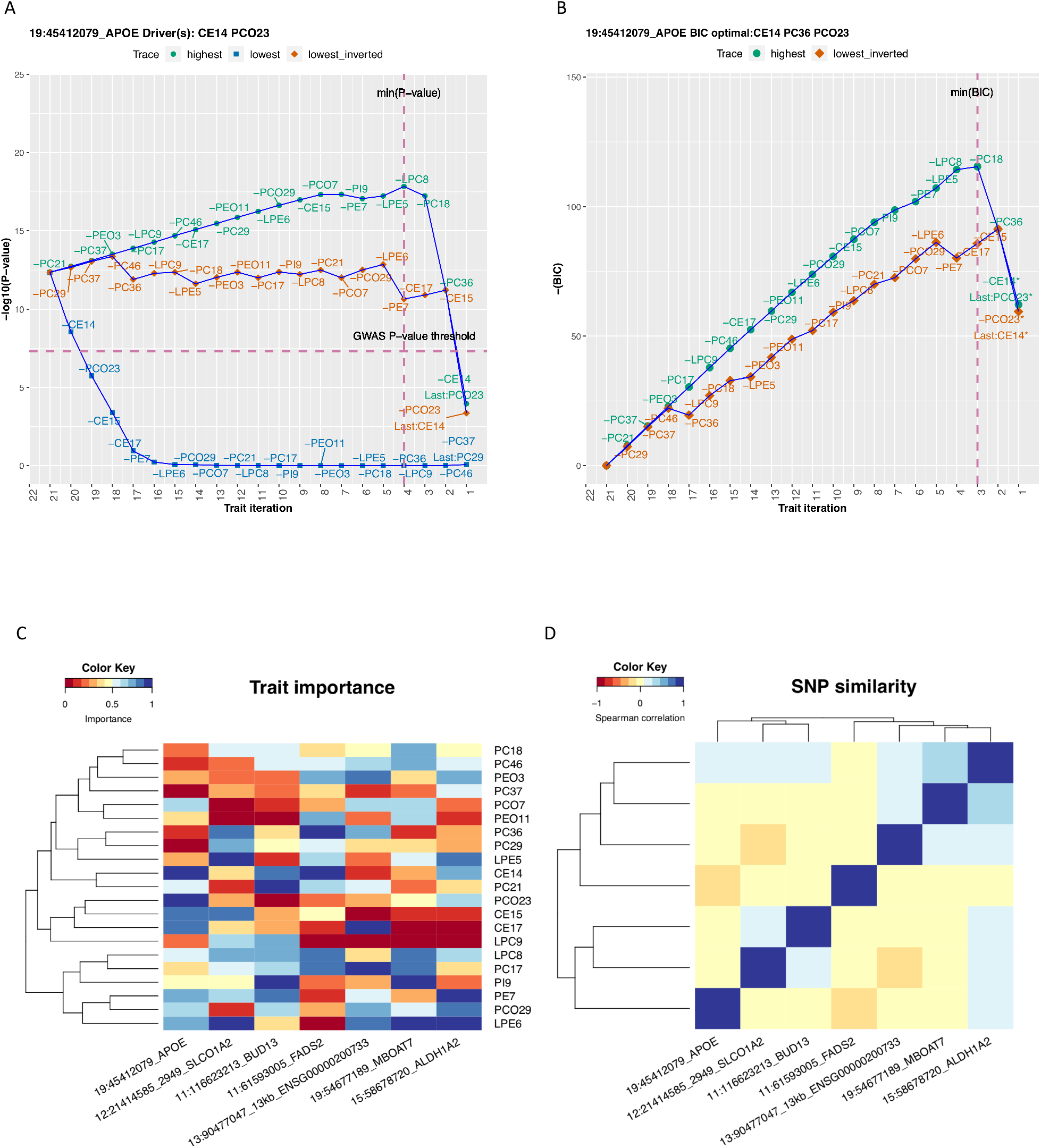
MetaPhat results using multivariate lipidomics data. A. Trace plot of rs7412 identifies CE14 and PCO23 as the traits driving the association. B, CE14, PC36, and PCO23 form the optimal subset as defined by minimum BIC (highest negative BIC). C. Trait importance map of each SNP is the rank on the lowest trace where the rankings are transformed to the range of 0 and 1 values, with darker blue shades representing the most important traits of the relevant lead association. D. SNP similarity based on the rank correlation on the lowest trace.

Variants rs66505542 near *BUD13* and rs261290 near *ALDH1A2* both have only one driver trait (*BUD13* PI9, *ALDH1A2* PE7) and 3 or 5 central traits (shown in Table 2), respectively, showing how MetaPhat has strongly reduced the multivariate association into much smaller groups of central traits. Earlier, the *APOA1* variant rs964184 within 100KB of rs66505542, has been reported associated with TG (lead trait, P = 7.0e-224), TC, HDL and LDL in GLGC data and rs66505542 itself with several cell phenotypes (platelet count, red cell distribution width, sum of eosinophil and basophil counts) in the GWAS catalogue, while rs261290 has been reported associated with HDL (lead trait, P = 1.0e-188), TC and TG in GLGC data (mapped to *LIPC* gene) and with HDL in the GWAS catalogue.

A very different picture emerges for rs174567 near *FADS1/2* as its 18 central traits shows its wide effects across the lipidomics traits studied here. Previously reported *FADS1/2* associations are with all lipid traits (TG lead trait, P = 7.0e-38) in GLGC data and with metabolite measurements and gallstones in the GWAS catalogue.

Trait importance map that clusters each variant based on the lowest trace is shown in Figure 2C and the similarity of the variants as measured by rank correlation of the traits on the lowest trace is shown in Figure 2D. The trace plots for the other six variants than rs7412 are shown in Figure S1.

## Validation and Global Lipids Genetics Consortium

We processed the Global Lipids Genetics Consortium (GLGC) GWAS study for four plasma lipids (HDL, LDL, TC and TG as listed in Table 1B). These correlated traits along with large sample sizes and available summary files are suitable for MetaPhat GWAS and decomposition. We focused on the 13 variants reported by GLGC to have associations with three or more lipid traits (Tables S2 and S3 from Willer et al., 2013). In Table 3, we validated that at 12 out of the 13 variants the same associations are confirmed by MetaPhat’s central traits. The only discordance was at rs6831256 (*DOK7*) where we found TC and TG as central traits compared to previously reported univariate associations with TC, TG and LDL. As TC and LDL are highly correlated, it is understandable that the smaller dimension of the set TC, TG may in some analyses be preferred over the set that also includes LDL. In Table S2, we further report high concordance between our central traits and GLGC variants found associated with two or more standard lipids.

**Table 3:** MetaPhat detection of driver and optimal lipid sets for 13 variants reported to be associated with at least three lipids by GLGC (12). We confirmed that the vast majority of the MetaPhat central traits are either the same or a subset of the reported GLGC associated lipids (11/13 for driver traits, 12/13 for BIC).

| Gene | Variant Chr:Pos | GLGC associated Lipids | GLGC Lead P-value | MetaPhat All traits P-value | MetaPhat Driver(s) | Without Driver(s) P-value | BIC Optimal set | Central traits |
| --- | --- | --- | --- | --- | --- | --- | --- | --- |
|  |  | **HDL lead** |  |  |  |  |  |  |
| *PIGV-NR0B2* | rs12748152 chr1:27138393 | HDL LDL TG | 1e-15 | 2.8e-23 | HDL LDL TG | 3.0e-06 | HDL LDL | HDL LDL TG |
| *PPP1R3B* | rs9987289 chr8:9183358 | HDL LDL TC | 2e-41 | 1.6e-76 | HDL TC LDL | 1.0e-04 | HDL LDL | HDL LDL TC |
| *LIPC (ALDH1A2)* | rs1532085 chr15:58683366 | HDL TC TG | 1e-188 | 0 | HDL TC TG | 6.4e-01 | HDL TC TG | HDL TC TG |
| *CETP* | rs3764261 chr16:56993324 | ALL | 1e-769 | 0 | ALL | NA | HDL LDL TG | ALL |
|  |  | **LDL lead** |  |  |  |  |  |  |
| *MIR148A* | rs4722551 7:25991826 | LDL TG TC | 4e-14 | 2.5e-24 | TG LDL TC | 2.0e-02 | LDL TG | LDL TG TC |
| *APOE* | rs4420638 19:45422946 | ALL | 2e-178 | 6.3e-210 | ALL | NA | LDL HDL TC | ALL |
|  |  | **TC Lead** |  |  |  |  |  |  |
| *TIMD4* | rs6882076 5:156390297 | TC LDL TG | 5e-41 | 1.3e-49 | TG TC LDL | 6.9e-01 | TC TG | TC LDL TG |
| *CILP2* | rs10401969 19:19407718 | TC TG LDL | 4e-77 | 1.3e-138 | TG TC LDL | 1.0e-01 | TC TG | TC TG LDL |
|  |  | **TG Lead** |  |  |  |  |  |  |
| *LRPAP1 (DOK7)* | rs6831256 4:3473139 | TG TC LDL | 2e-12 | 6.3e-16 | TG TC | 1.0e-07 | TG TC | TG TC |
| *ANGPTL3* | rs2131925 1:63025942 | TG LDL TC | 3e-74 | 7.8e-157 | TG LDL TC | 9.5e-05 | TG TC HDL | ALL |
| *TRIB1* | rs2954029 8:126490972 | ALL | 1e-107 | 1.6e-148 | ALL | NA | TG TC LDL | ALL |
| *FADS123* | rs174546 11:61569830 | ALL | 7e-38 | 1.3e-104 | ALL | NA | ALL | ALL |
| *APOA1* | rs964184 11:116648917 | ALL | 7e-224 | 7.9e-264 | ALL | NA | TG TC | ALL |

## Performance

For computing the test statistic, MetaPhat uses metaCCA that, for a single SNP, has previously been shown to reliably estimate the results of standard CCA applied to individual level data (canoncorr function in Matlab) (Cichonska et al., 2016). Additionally, we also empirically validated MetaPhat multivariate findings with GLGC results.

MetaPhat considerably cuts down the computational demands of comprehensive subset testing. With K traits, there are 2^K^-1non-empty subsets which quickly become infeasible to systematically assess, while MetaPhat only considers about K^2^ models. For example, in our example with K=21 traits, the gain in performance is about 4,700-fold compared to the complete subset testing. To further increase performance and usability, we have implemented flexibility for multi-thread processing to enable high performance and memory efficiency. On a moderate Google cloud image (16 vCPUs, 8 GB), the complete MetaPhat workflow for our lipidomics analysis, containing 21 lipids and 8.5 million SNPs, completed in less than 2.5 hours (143 minutes). Using 10 processors and 9 gigabytes of memory, the GLGC job with the 4 basic lipids and 2.4 million imputed SNPs completed in 24 minutes. MetaPhat also allows decomposition and plotting of custom SNPs. For example, the custom analysis of the 13 GLGC variants associated with 3 or more traits, shown in Table 3, was reran on existing GLGC MetaPhat results and decomposition and plotting took only 2 minutes. We note that the run time could be longer on shared servers but also substantially shorter using more powerful dedicated cloud images.

## Discussion

It is expected that a particular genetic variant may affect only a subset of related biomarkers that are risk factors of complex disorders such as T2D or coronary heart disease. We implemented MetaPhat to systematically decompose and visualize statistically significant multivariate genome-phenome associations into a smaller group of central traits, based only on univariate GWAS summary statistics. We are not aware of comparable software to MetaPhat that would automatically carry out multivariate GWAS and identify central traits for the associations from summary statistics. ASSET (Bhattacharjee et al., 2012) aims to find the best trait subsets within a pool of multiple studies and has been applied particularly for case-control studies. MTAG (Turley et al., 2018) can be applied to GWAS results of multiple related traits and overlapping samples but its aim is to improve the accuracy of the univariate effect sizes by using the information from correlated traits rather than decomposing the multivariate association to individual traits.

In our results from analysis of 21 lipidomics traits, we demonstrated that the *APOE* association benefited from multivariate testing (driven by CE14 and PCO23 traits) as the univariate P-value was insignificant (P > 1e-4) across all 21 GWAS traits but multivariate P-value was low (P < 5e-13). Additionally, MetaPhat decomposed most variants to substantially smaller sets of central traits than 21. On the other hand, the essential role of *FADS2* gene region in regulating unsaturation in fatty acids was clearly reflected in MetaPhat results as we observed as many as 18 central traits at the lead variant. Provided that the exact mechanistic roles of polyunsaturated lipids towards heart disease (Malovini et al., 2016, Pizzini et al., 2017, Teslovich et al., 2010) are under active investigation, our findings warrant further evaluation. We further confirmed good concordance (60/67, Table S2) with MetaPhat central traits with respect to the earlier reported GLGC associations with two or more standard lipids, and excellent concordance (12/13) with the associations with three or more standard lipids.

MetaPhat optimal subsets are derived from the minimum BIC score representing the model that best describes the data when we account for both the model fit and the model dimension. Qualitatively BIC statistic is similar to the widely-used AIC (Akaike, 1973) statistic but quantitatively BIC differs from AIC by favoring smaller dimensions which also improves the interpretation of the optimal models. As intuitively expected, and as seen in Table 2, the driver traits tend to be members of the optimal set although they do not always agree, since the driver traits are defined by a GWAS-specific criterion of P-value threshold 5e-8 which does not need to coincide with the optimal subset chosen by a more statistically justified BIC criterion.

Our software implements flexible parameters for custom multi-thread chunking to enable high performance, genome-wide, multi-trait meta-analysis while integrating metaCCA for multivariate testing followed by systematic decomposition of traits. Thus, a limitation of MetaPhat is that it relies on metaCCA but other multivariate GWAS algorithms could also be used provided that these methods can work with univariate GWAS results as inputs and produce suitable metrics that can be used to derive the model comparison statistics. With regard to false positives, we are using the standard GWAS cutoff (P = 5e-8), as we are doing only a single multivariate GWAS to pick the lead variants. This cutoff can be adjusted according to the preferences of the users. MetaPhat also optionally allows running metaCCA+ (Cichonska et al., 2016), shown to protect against false positives via shrinkage that adds robustness to the analysis.

Finally, we remind that MetaPhat decompositions are sequential, dropping one trait at a time, and hence are not guaranteed to produce the globally optimal subset. Additionally, for highly correlated traits, such as LDL and total cholesterol, the choice of which one is dropped first may not be completely robust to small changes in data.

The ability of MetaPhat to identify and visualize central traits will also be valuable in supporting efforts and pipelines (Fatumo et al., 2019) comparing results between univariate and multivariate associations as well as in studies that aim to increase specificity of multi-trait associations. We also expect that the multi-phenotype clustering results of MetaPhat can assist researchers investigating disease subtypes.

## Availability and requirements

Project home page: https://sourceforge.net/projects/meta-pheno-association-tracer/

Operatingsystem: Independent

Languages: Python 2.7+, Python 3.5+, R 3.4

Licence: MIT

## Funding

This work was supported by the Academy of Finland (grants no. 288509, 312076, 319181, 325999).

## Conflicts of Interest

none declared.

## Notes

#### Summary of Updates

The manuscript has been expanded from applications note to full methods paper. The revision includes BIC optimal set implementation, expanded lipidomics results, workflow and validation using Global Lipids Genetics Consortium summaries.

https://sourceforge.net/projects/meta-pheno-association-tracer/

