## Supplement data for "MetaPhat: Detecting and decomposing multivariate associations from univariate genome-wide association statistics"

### Supplemental data

A

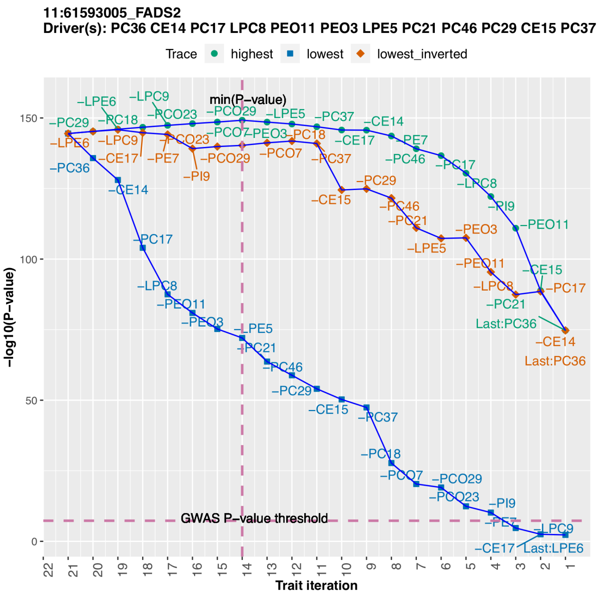


B

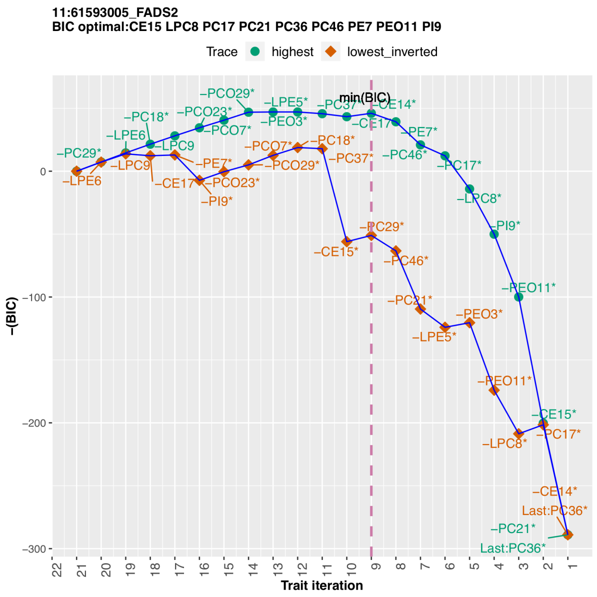


C

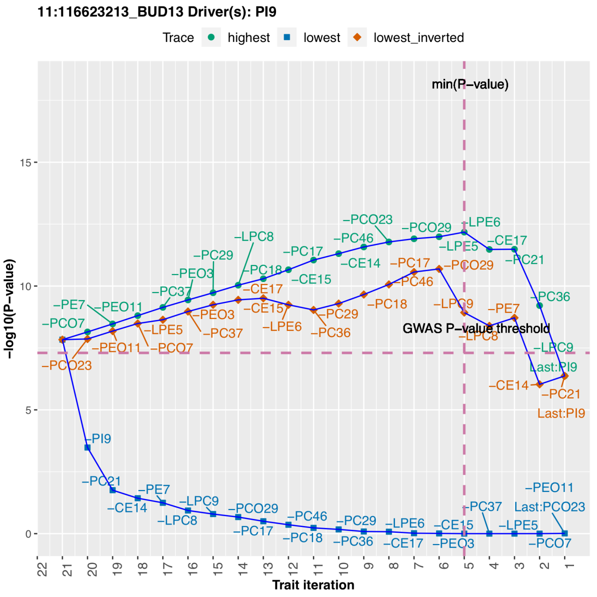


D

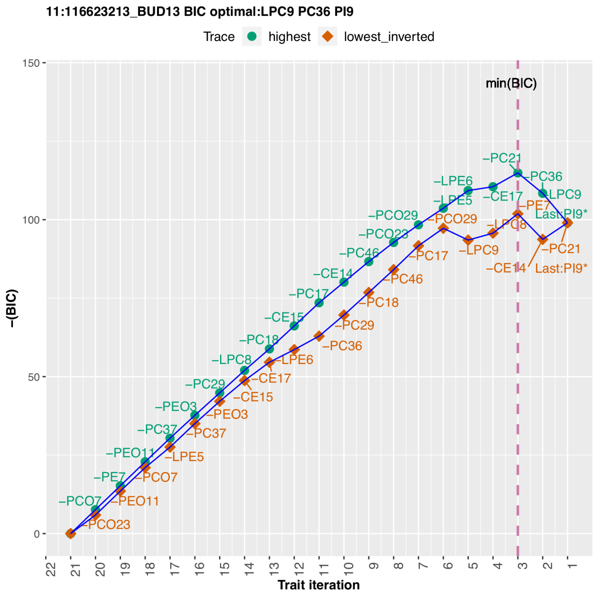

E

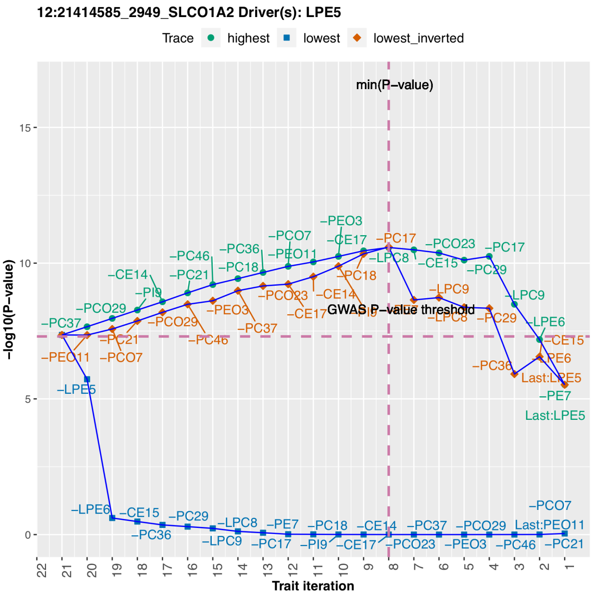
12:21414585_2949_SLCO1A2 Driver(s): LPE5
Trace highest lowest lowest_inverted
min(P−value)
GWAS P−value threshold
−log10(P−value)
Trait iteration

F

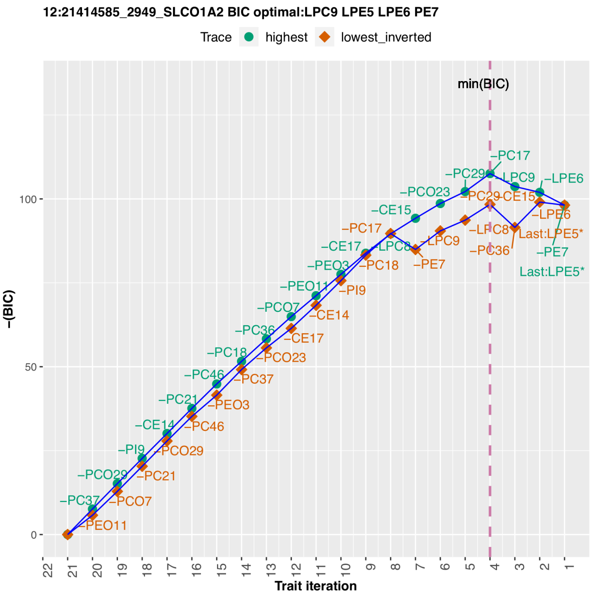
12:21414585_2949_SLCO1A2 BIC optimal:LPC9 LPE5 LPE6 PE7
Trace highest lowest_inverted
min(BIC)
−(BIC)
Trait iteration

G

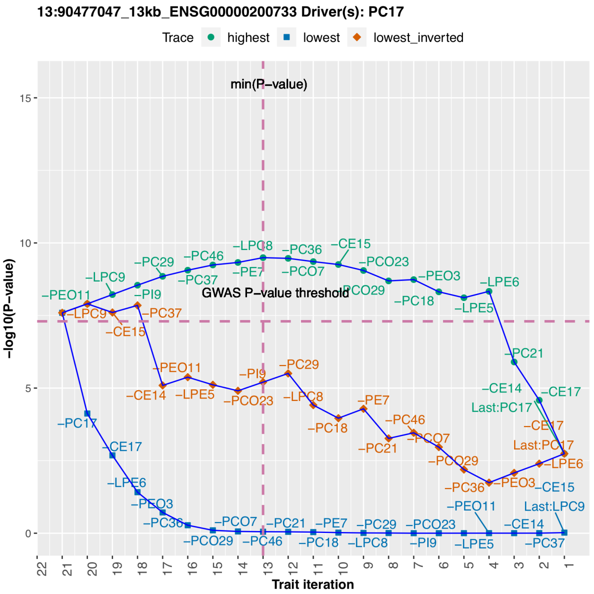
13:90477047_13kb_ENSG00000200733 Driver(s): PC17
Trace highest lowest lowest_inverted
min(P−value)
GWAS P−value threshold
−log10(P−value)
Trait iteration

H

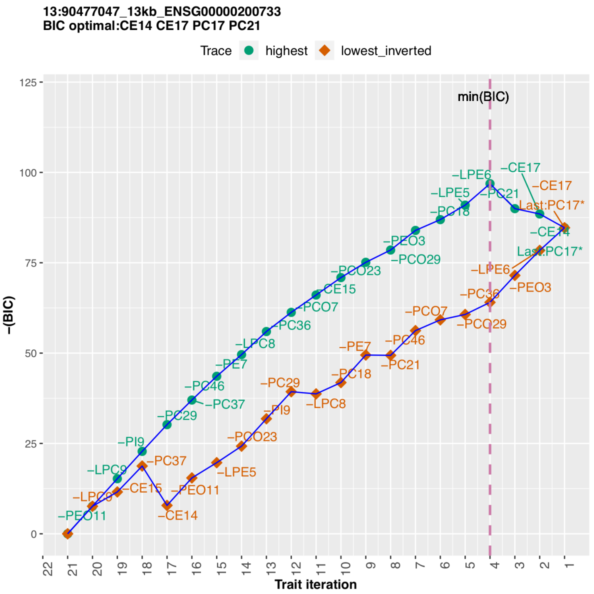
13:90477047_13kb_ENSG00000200733
BIC optimal:CE14 CE17 PC17 PC21
Trace highest lowest_inverted
min(BIC)
−(BIC)
Trait iteration

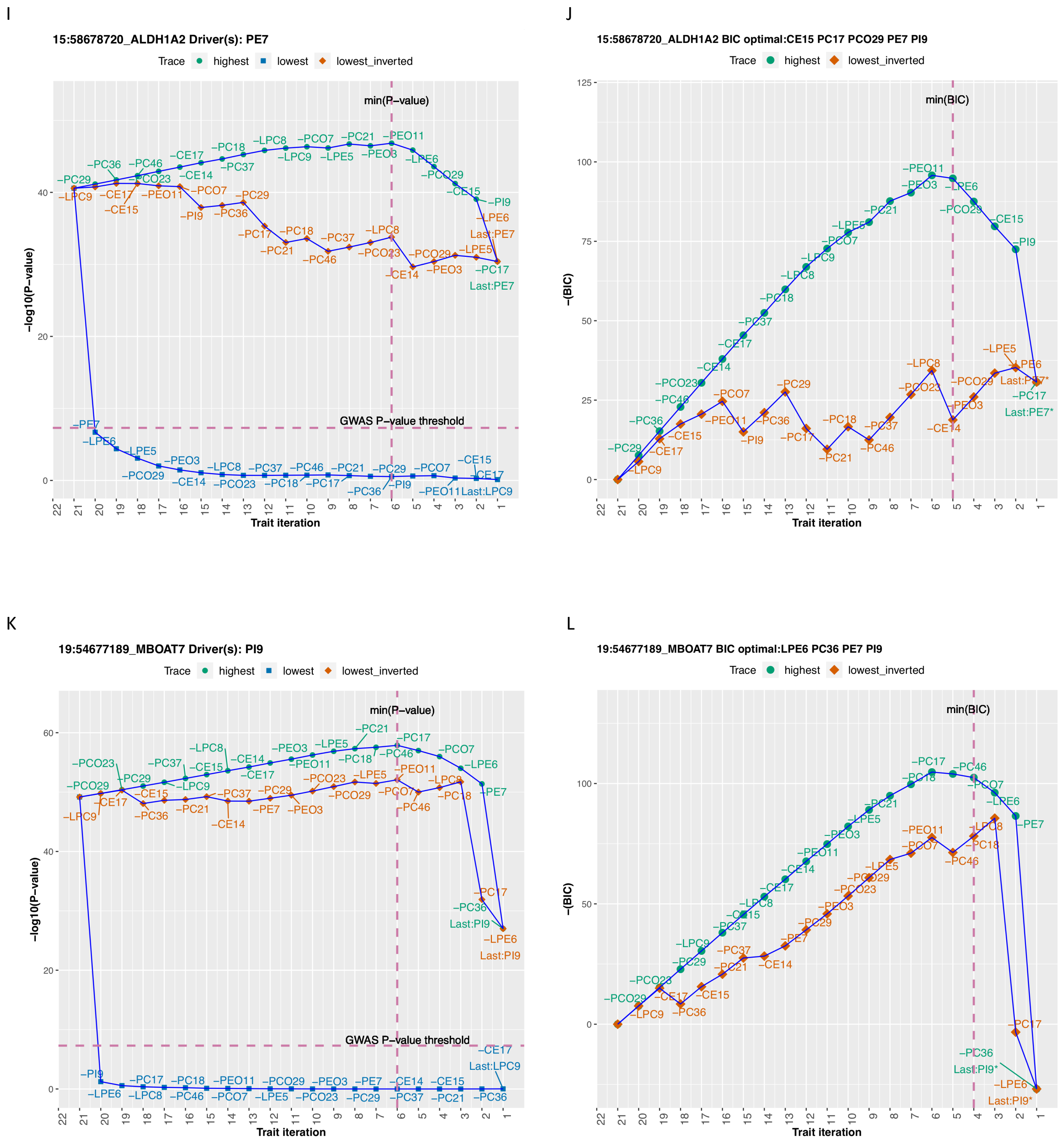


Figure S1 – P-value and BIC trace plots of lead SNP associations detected by MetaPhat decomposition. Additional details are listed in Table 2 of the main text.

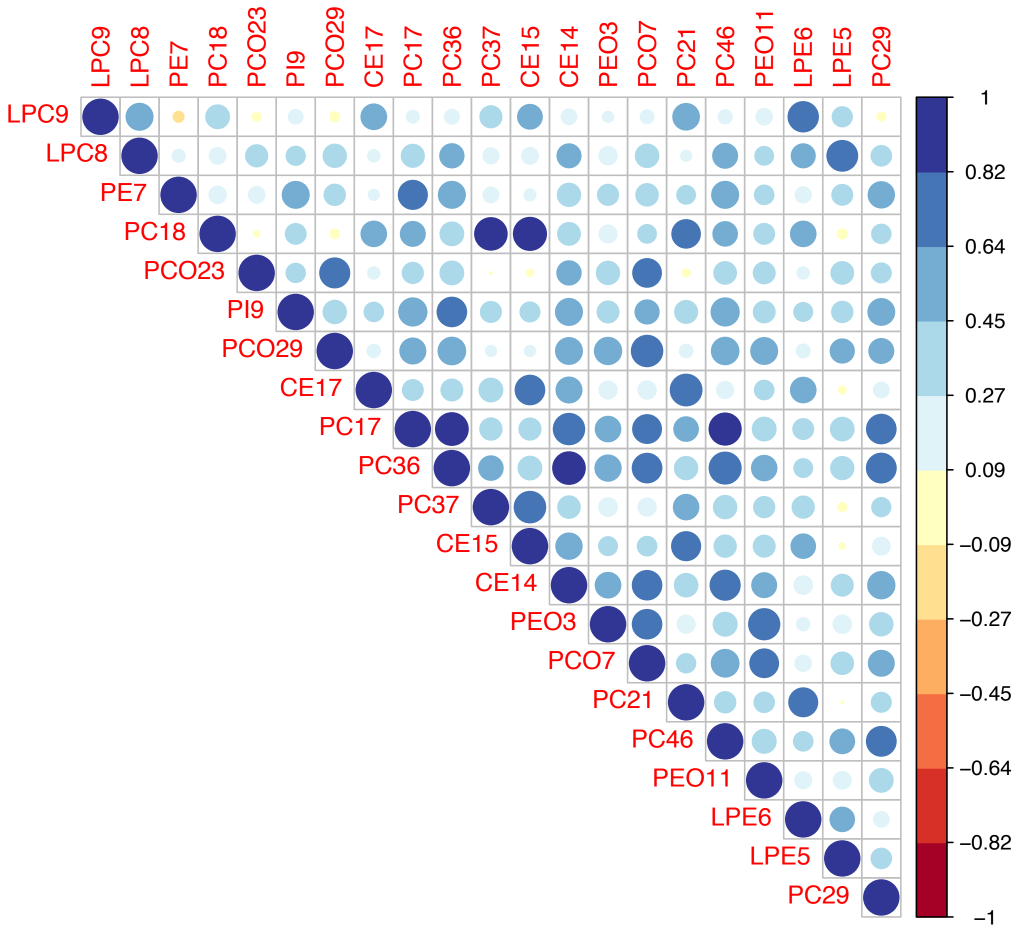


**Figure S2:** Correlation map of 21 heritable lipid traits computed from univariate beta coefficients using metaCCA.

Table S1 – MetaPhat detection of driver and optimal lipids for 69 variants reported to associate with two or more essential GLGC lipids (Willer et al., 2013). As two variants[1] are not included in the summary statistics, the list of 67 variants are group by the reported leading trait, and listed with their central traits. We confirmed that 60/67 of the associated GLGC lipids were either identical or a subset of the central traits.

| **Gene** | **VariantId Chr:Pos** | **GLGC associated Lipids** | **GLGC Lead P-value** | **MetaPhat All traits P-value** | **MetaPhat Driver(s)** | **Driver(s) Dropped P-value** | **BIC Optimal set** | **Central traits** |
|---|---|---|---|---|---|---|---|---|
| | | **HDL lead** | | | | | | |
| PIGV-NR0B2 | rs731839 chr1:27138393 | HDL LDL TG | 1e-15 | 2.8e-23 | HDL LDL TG | 3.0e-06 | HDL LDL | HDL LDL TG |
| RSPO3 | rs1936800 chr6:127436064 | HDL TG | 3e-10 | 6.6e-09 | TG | 7.9e-08 | HDL | HDL TG |
| MARCH8-ALOX5 | rs970548 10:46013277 | HDL TC | 2e-10 | 1.8e-17 | HDL TC | 9.1e-04 | HDL TC | HDL TC |
| FTO | rs1121980 chr16:53809247 | HDL TG | 7e-09 | 4.9e-09 | TG | 1.6e-07 | HDL | HDL TG |
| GALNT2 | rs4846914 chr1:230295691 | HDL TG | 4e-41 | 5.3e-51 | HDL TG | 1.1e-04 | HDL TG | HDL TG |

| | | | | | | | | |
|---|---|---|---|---|---|---|---|---|
| IRS1 | rs2972146 chr2:227100698 | HDL TG | 2e-17 | 4.6e-22 | HDL TG | 4.6e-01 | HDL TG | HDL TG |
| PPP1R3B | rs9987289 chr8:9183358 | HDL LDL TC | 2e-41 | 1.6e-76 | HDL TC LDL | 1.0e-04 | HDL LDL | ALL |
| TTC39B | rs581080 chr9:15305378 | HDL TC | 1e-19 | 6.9e-46 | HDL TC | 5.6e-08 | HDL LDL TG | ALL |
| ABCA1 | rs1883025 chr9:107664301 | HDL TC | 2e-65 | 1.9e-131 | HDL TC TG | 2.9e-07 | HDL LDL TG | ALL |
| ZNF664 | rs4765127 chr12:124460167 | HDL TG | 8e-10 | 1.2e-12 | HDL TG | 1.6e-03 | HDL TC | HDL TC TG |
| LIPC (ALDH1A2) | rs1532085 chr15:58683366 | HDL TC TG | 1e-188 | 0 | HDL TC LDL | 6.4e-01 | HDL TG LDL | ALL |
| CETP | rs3764261 chr16:56993324 | ALL | 1e-769 | 0 | ALL | 1.6e-25 | HDL LDL TG | ALL |
| LIPG | rs7241918 chr18:47160953 | HDL TC | 1e-44 | 5.4e-59 | HDL TC | 9.5e-04 | HDL LDL TG | ALL |
| HNF4A | rs1800961 chr20:43042364 | HDL TC | 2e-34 | 1.5e-60 | HDL TC LDL | 7.1e-01 | HDL LDL TG | ALL |
| PLTP (ZNF335) | rs6065906 chr20:44554015 | HDL TG | 5e-40 | 8.9e-50 | HDL TG | 1.3e-03 | HDL TG | HDL TG |
| | | **LDL lead** | | | | | | |
| INSIG2 | rs10490626 2:118835841 | LDL TC | 2e-12 | 9.8e-14 | TG LDL HDL | 5.3e-02 | TG LDL | LDL TG |
| LOC84931 | rs2030746 2:121309488 | LDL TC | 9e-09 | 5.9e-07 | NA | NA | LDL | LDL |
| CMTM6 | rs7640978 3:32533010 | LDL TC | 1e-08 | 1.8e-08 | LDL | 1.4e-07 | LDL | LDL |
| ACAD11 (DNAJC13) | rs17404153 chr3:132163200 | LDL HDL | 2e-09 | 1.8e-08 | LDL | 1.2e-07 | LDL | LDL |
| CSNK1G3 | rs4530754 5:122855416 | LDL TC | 4e-12 | 1.7e-12 | LDL TC | 7.9e-01 | LDL | LDL TC |
| MIR148A | rs4722551 7:25991826 | LDL TG TC | 4e-14 | 2.5e-24 | TG LDL TC | 2.0e-02 | LDL TG | LDL TG TC |
| SOX17 | rs10102164 8:55421614 | LDL TC | 4e-11 | 5.0e-12 | LDL TC | 1.0e-02 | LDL | LDL TC |
| PCSK9 | rs2479409 1:55504650 | LDL TC | 3e-50 | 3.2e-53 | LDL TC | 2.8 e-02 | LDL | LDL TC |
| SORT1 | rs629301 1:109818306 | LDL TC | 5e-241 | 1.3e-270 | LDL TC TG | 6.3e-01 | LDL TG | LDL TC TG |
| APOB | rs1367117 2:21263900 | ALL | 1e-182 | 5.0e-197 | ALL | 7.9e-09 | LDL TC | ALL |
| ABCG5 | rs4299376 2:44072576 | LDL TC | 4e-72 | 1.0e-79 | LDL TC | 6.3e-03 | LDL TC | LDL TC |
| MYLIP | rs3757354 6:16127407 | LDL TC | 1e-17 | 1.3e-17 | LDL TC | 6.3e-01 | LDL | LDL TC |
| HFE | rs1800562 6:26093141 | LDL TC | 8e-14 | 1.6e-15 | TG LDL TC | 2.4e-01 | LDL | LDL TG TC |
| LPA | rs1564348 6:160578860 | LDL TC | 3e-21 | 1.3e-23 | LDL TC | 1.2e-03 | TC | LDL TC |
| PLEC1 | rs11136341 8:145043543 | LDL TC | 7e-12 | 1.0e-11 | LDL TC | 6.3e-01 | LDL | LDL TC |
| ST3GAL4 | rs11220462 11:126243952 | LDL TC | 7e-21 | 6.3e-22 | LDL TC | 7.9e-04 | LDL | LDL TC |
| OSBPL7 | rs7206971 17:45425115 | LDL TC | 1e-07 | 1e-09 | HDL TG LDL | 3.2e-07 | LDL HDL | ALL |
| LDLR | rs6511720 19:11202306 | LDL TC | 4e-262 | 5.0e-296 | LDL TC | 6.3e-05 | LDL TG | LDL TC TG |

| | | | | | | | | |
|---|---|---|---|---|---|---|---|---|
| APOE (APOC) | rs4420638 19:45422946 | LDL TC HDL | 2e-178 | 6.3e-210 | ALL | 1.6e-14 | LDL HDL TC | ALL |
| TOP1 | rs6029526 20:39672618 | LDL TC | 5e-18 | 1.0e-15 | LDL TC | 1.1e-01 | LDL | LDL TC |
| | | **TC Lead** | | | | | | |
| UGT1A1 | rs11563251 2:234679384 | TC LDL | 6e-09 | 2.0e-08 | TC LDL | 5.0e-02 | TC | TC LDL |
| VLDLR | rs3780181 9:2640759 | TC LDL | 7e-10 | 6.3e-10 | TC LDL | 5.1e-01 | TC | TC LDL |
| DLG4 | rs314253 17:7091650 | TC LDL | 3e-10 | 1.6e-09 | LDL TC | 3.9e-02 | LDL | TC LDL |
| PPARA | rs4253772 22:46627603 | TC LDL | 1e-08 | 1.9e-08 | TG | 3.2e-07 | LDL | LDL TG |
| LDLRAP1 | rs12027135 1:25775733 | TC LDL | 5e-12 | 4.0e-14 | TC LDL | 6.3e-02 | LDL | TC LDL |
| MOSC1 (MARC1) | rs2642442 1:220973563 | TC LDL | 3e-11 | 2.5e-17 | LDL TC | 1.3e-05 | HDL LDL | TC LDL HDL |
| IRF2BP2 | rs514230 1:234858597 | TC LDL | 5e-14 | 1.3e-11 | TC LDL | 1.6e-01 | TC | TC LDL |
| HMGCR | rs12916 5:74656539 | TC LDL | 5e-74 | 1.6e-90 | TC LDL | 2.0e-01 | TC LDL | TC LDL |
| TIMD4 | rs6882076 5:156390297 | TC LDL TG | 5e-41 | 1.3e-49 | TG TC LDL | 6.9e-01 | HDL LDL TG | ALL |
| HLA | rs3177928 6:32412435 | TC LDL | 1e-21 | 2.0e-24 | HDL TC LDL | 3.2e-03 | HDL LDL TG | ALL |
| FRK | rs9488822 6:116312893 | TC LDL | 1e-09 | 1.3e-10 | TG TC LDL | 2.0e-01 | LDL | TC TG LDL |
| DNAH11 | rs12670798 7:21607352 | TC LDL | 1e-16 | 6.3e-17 | TC LDL | 2.5e-02 | TC | TC LDL |
| NPC1L1 | rs2072183 7:44579180 | TC LDL | 4e-15 | 7.9e-16 | TC LDL | 4.0e-03 | LDL | TC LDL |
| CYP7A1 | rs2081687 8:59388565 | TC LDL | 9e-12 | 5.0e-13 | TG TC LDL | 5.0e-01 | TC | TC TG LDL |
| GPAM | rs2255141 10:113933886 | TC LDL | 7e-16 | 1.0e-36 | ALL | 1.3e-13 | HDL LDL | ALL |
| BRAP | rs11065987 12:112072424 | TC LDL | 2e-16 | 7.9e-24 | TC LDL TG | 2.0e-01 | LDL HDL | ALL |
| HNF1A | rs1169288 12:121416650 | TC LDL | 4e-17 | 2.5e-23 | LDL TC | 4.0e-03 | LDL HDL | TC LDL HDL |
| HPR | rs2000999 16:72108093 | TC LDL | 7e-41 | 1.6e-49 | LDL TC | 1.3e-06 | LDL TC | TC LDL |
| CILP2 | rs10401969 19:19407718 | TC LDL TG | 4e-77 | 1.3e-138 | TG HDL TC | 1.0e-01 | HDL LDL TG | ALL |
| MAFB | rs2902940 20:39091487 | TC LDL | 9e-10 | 2.5e-11 | TC LDL | 1.8e-02 | LDL | TC LDL |
| | | **TG Lead** | | | | | | |
| LRPAP1 (DOK7) | rs6831256 4:3473139 | TG TC LDL | 2e-12 | 6.3e-16 | TG TC | 1.0e-07 | TG TC | TG TC |
| VEGFA | rs998584 6:43757896 | TG HDL | 3e-15 | 1.2e-16 | TG HDL | 5.3e-01 | HDL | TG HDL |
| PEPD | rs731839 19:33899065 | TG HDL | 3e-09 | 2.5 e-11 | TG HDL | 1.3e-01 | TG | TG HDL |
| ANGPTL3 | rs2131925 1:63025942 | TG LDL TC | 3e-74 | 7.8e-157 | TG TC LDL | 9.5e-05 | TG TC HDL | TG LDL TC |
| GCKR | rs1260326 2:27730940 | TG TC | 2e-239 | 2.0e-290 | TG TC LDL | 1.7e-03 | TG TC HDL | TG TC LDL |
| MLXIPL | rs17145738 7:72982874 | TG HDL | 9e-99 | 7.9e-105 | TG HDL | 7.8e-06 | TG | TG HDL |

| | | | | | | | | |
|---|---|---|---|---|---|---|---|---|
| NAT2 | rs1495741 8:18272881 | TG TC | 3e-12 | 5.9e-14 | TG | 1.1e-07 | TG | TG |
| LPL | rs12678919 8:19844222 | TG HDL | 2e-199 | 1.0e-253 | TG HDL | 9.5e-02 | TG LDL HDL | TG HDL LDL |
| TRIB1 | rs2954029 8:126490972 | ALL | 1e-107 | 1.6e-148 | ALL | NA | TG LDL | ALL |
| FADS123 | rs174546 11:61569830 | ALL | 7e-38 | 1.3e-104 | ALL | NA | ALL | ALL |
| APOA1 | rs964184 11:116648917 | ALL | 7e-224 | 7.9e-264 | ALL | NA | TG TC | ALL |
| LRP1 | rs11613352 12:57792580 | TG HDL | 9e-14 | 7.9e-16 | TG HDL | 1.6e-04 | TG HDL | TG HDL |

Two associations not found in lipid summaries: ABO (rs7941030) LDL TC and UBASH3B (rs7941030):TC HDL

**Table S2**: The input switches of MetaPhat are shown with the --help option. Table below outlines the argument switches and their default values.

| Switches | Description and default value |
|---|---|
| --help | Shows all arguments and lists relevant default values |
| --phenotypes | Required, parameter is a file that defines GWAS summary files for testing (key:full_path)<br><br>One summary file per line, in this format<br>pheno1:/path/summary1.gz<br>pheno2:/path/summmary2.gz<br>...<br>See example:<br>https://sourceforge.net/projects/meta-pheno-association-tracer/files/test_inputs/global_lipid_summaries2 |
| --outlabel | Required, this defines a label for output files and plotting, must be alphanumeric and without spaces. |
| --nsamples | Required, enter in estimate number of subjects the GWAS summaries were based on. |
| --nsamples_dev | Optional, this defines the allowed deviating percentage of missing samples for variants to be included in analysis. It only applies if N is defined in GWAS summaries. Defaults to .25 (25%, for example, if sample size N =1000, variants with <750 or >1250 will be excluded) |
| --chunksize | Required, this sets the number of SNPs processed by metaCCA on each batch. Defaults to 100000 |
| --parallel | Required, this defines the number of threads, or batch jobs executed at the same time. The entry depends on your processor and whether it is shared. Defaults to 5. |
| --waittime | Required, this defines the number of seconds each batch submission approximately runs. Waittime is important and depends on the chunksize and parallel threads, our testing is that this should be around 120-180 seconds for chunks of 100000 and 6-8 threads on 16Gb 16Core nonshared environment. On shared server, we recommend using higher wait time. Defaults to 180. |
| --r1 | Optional, this defines the output column from metaCCA to sort by during clumping. metaCCA outputs r1 and pvalue and we recommend r1 as pvalue can be INF and problematic for true sort of importance. Defaults to 1 meaning CCA correlation. |
| --outdir | Required, this defines the complete path to store outputs. |
| --gwascutoff | Required. This defines the pvalue cutoff, -1(math.log(pv, 10)) Defaults to 7.3, ~0.00000001 approximating recommended GWAS threshold |

| | |
|---|---|
| --grch | Optional, this defines the human ENSEMBL (Hunt et al. 2018) build version, possible values are 37 and 38. Defaults to 37. |
| --exclude | Optional, this defines a file where variants can be excluded even if the variant is significant. One variant per line and see example: https://sourceforge.net/projects/meta-pheno-association-tracer/files/test_inputs/exclude.dat |
| --interested | Optional, this defines a file where variants are included in the trace results even if variant pvalue does not pass cutoff. One variant per line and see example: https://sourceforge.net/projects/meta-pheno-association-tracer/files/test_inputs/interested.list |
| --maf_range | Optional, this defines the variant frequency min:max, range to include. Example values can be .01:.5, meaning maf values >.5 and variants <.01 will be excluded. Only applies if all input GWAS summaries include this field, and the field column header needs to be effect_allele_frequency or maf. For possible guidelines, see UK Biobank (item 2): http://www.nealelab.is/blog/2017/9/11/details-and-considerations-of-the-uk-biobank-gwas |
| --clump | Required, this defines the base window to sort variants and their pvalues/r1 canonical correlation. Defaults to 500000. |
| --neglogval | Optional. Parameter defines the maximal neg log pvalue for INF values. Defaults to 400. |
| --plot_intermediate | Optional. Parameter tells program to plot intermediate lead cca value for all iterations. Defaults to 0. |
| --title_font | Optional. Parameter defines the font size of title in trace plot. Defaults to 12. |
| --element_font | Optional. Parameter defines the font size of element label in trace plot. Defaults to 5. |
| --axis_font | Optional. Parameter defines the font size of axis label in trace plot. Defaults to 12. |
| --legend_font | Optional. Parameter defines the font size of legend in trace plot. Defaults to 12. |
| --Rscript | Required, this defines the full path to Rscript executable. Defaults to /usr/bin/Rscript |
